# Comprehensive mouse gut metagenome catalog reveals major difference to the human counterpart

**DOI:** 10.1101/2021.03.18.435958

**Authors:** Silas Kieser, Evgeny M. Zdobnov, Mirko Trajkovski

## Abstract

Mouse is the most used model for studying the impact of microbiota on its host, but the repertoire of species from the mouse gut microbiome remains largely unknown. Here, we construct a Comprehensive Mouse Gut Metagenome (CMGM) catalog by assembling all currently available mouse gut metagenomes and combining them with published reference and metagenome-assembled genomes. The 50’011 genomes cluster into 1’699 species, of which 78.1% are uncultured, and we discovered 226 new genera, 7 new families, and 1 new order. Rarefaction analysis indicates comprehensive sampling of the species from the mouse gut. CMGM enables an unprecedented coverage of the mouse gut microbiome exceeding 90%. Comparing CMGM to the human gut microbiota shows an overlap 64% at the genus, but only 16% at the species level, demonstrating that human and mouse gut microbiota are largely distinct.

## Introduction

Mouse is the most used model for studying the microbiota importance due to several factors: availability of samples from different parts of the gastrointestinal tract, treatment options, controlled housing environment and diet, defined genetic background, and ethical considerations. However, the mouse gut microbiota has been poorly characterized, and only a fraction of the diversity observed by 16S rDNA sequencing is represented by genomes in public databases (Lagkouvardos et al., 2016). The majority of mouse microbiome studies are performed by sequencing variable regions of the 16S, sometimes mislabeled as metagenomics. While this technique has allowed a general overview of the microbiota down to the genus level, it is not suited for identifying species for most organisms (Johnson et al., 2019). Different species from the same genus and even subspecies from the same species can exert distinct functions (Costea et al., 2017), stressing the importance of annotating the gene content at the lowest taxonomic level.

Shotgun metagenomics allows studying the full microbiota diversity of an environment, including uncultured microorganisms, viruses, and plasmids. However, its interpretation is limited by the availability of reference genomes. Previous efforts led to the creation of a gene catalog of the mouse metagenome (MGC v1) (Xiao et al., 2015), by sequencing fecal samples from mice with different genotypes and housed in different conditions. This catalog enables the functional annotation of genes and allows up to a 50% mapping rate of fecal shotgun sequences. However, the mapping rate of sequences from cecum samples is only 37%, and the catalog does not contain genomic references. Recently developed algorithms enable the assembly of genomes from metagenomes, leading to a recovery of new species from the human gut and other environments (Almeida et al., 2019; Nayfach et al., 2019; Parks et al., 2017; Pasolli et al.; Stewart et al., 2018). The integrated mouse gut metagenomic catalog (iMGMC) (Lesker et al., 2020) increased the fraction of reads mapped to genes compared to the MGC v1. However, mapping to the recovered metagenome-assembled genomes (MAGs) remained about 40% (Lesker et al., 2020). Lesker *et al.* also generated a set of 13,619 mouse-specific MAGs (mMAG) not integrated into the iMGMC, which was made available for further studies.

Here we report the creation of the Comprehensive Mouse Gut Metagenome (CMGM) collection by assembling gut microbiomes sequenced by us and all publicly available mouse metagenomes. This resource improves the mapping rate of genomic reads from mouse fecal and cecum metagenomes to over 90.5%, provides full classification down to species level, and enables uncovering compelling functional insights linking them to the driver species. This nearly complete catalog of the mouse gut bacterial species allows comparison between the newly assembled mouse gut microbiomes and the human counterpart, highlighting major differences between human and mouse.

## Results

### Assembly of high-quality genomes from mouse gut metagenomes

We selected all metagenomic datasets associated with the mouse intestinal tract that are sequenced as paired-ends from the NCBI sequence read archive. To these, we added samples generated by our lab resulting in 1226 datasets (Table S1). Each sample was processed using metagenome-atlas (Kieser et al., 2020), which handles pre-processing, assembly, and binning of the metagenome datasets. We included all mouse-associated bacterial genomes retrieved from RefSeq belonging to 331 species (Table S2, Fig. S1), which also incorporates genomes from mouse specific culture collections: Oligo-mouse-microbiota (Garzetti et al., 2017) (12 genomes), and Mouse Gut Microbial Biobank (mGMB, 41 genomes) (Liu et al., 2020). As genomes of the mouse Intestinal Bacterial Collection (miBC, 53 genomes) (Lagkouvardos et al., 2016) were not available, we assembled them from the raw reads. To this extensive new set, we included all MAGs produced by Lesker *et al*., thereby adding comprehensiveness to the newly assembled catalog (Fig. S1). All genomes were filtered based on fragmentation (N50 >5000) and a quality score calculated from the output of checkM (Parks et al., 2015) as ‘completeness minus 5 times contamination’. Bins with a quality score of <50 were excluded, resulting in a set of 49,’195 MAGs of which 15’355 (31%) had high quality (Quality score >90, Fig. S2A+B). Surprisingly, some reference genomes had contamination values of 100%, suggesting that the sequenced genomes consist of multiple strains. In total, 13 reference genomes did not pass the quality filtering, and we included 816 reference genomes in the CMGM collection, resulting in a total of 50’011 genomes.

While MAGs were more fragmented and had a lower median quality score than the reference genomes, the quality score and N50 of the high-quality MAGs were comparable to the values of the references (Fig. S2B). For 60% of the reference genomes, we recovered MAGs that align to them with high coverage and identity (average nucleotide identity (ANI) >95%, IQR 94-99%, Fig. S2C). This result validates our metagenome assembly approach to recover “reference quality” genomes *de novo*. Some of the remaining differences might be attributed to strain variation, as the coverage is higher for more similar genomes (Fig. S2C).

Since we assembled genomes from individual samples, the same strain could have been recovered multiple times, especially if different gut locations of the same mouse were sampled. To remove this potential redundancy, we clustered the genomes based on the ANI calculated using bindash (Zhao, 2018). 95% ANI was used as a threshold to delineate genomes from the same species (Jain et al., 2018; Olm et al., 2019). For each species cluster, the genome with the highest quality and lowest fragmentation was selected as representative, but reference genomes were preferred over MAGs. The species representatives were annotated using DRAM (Shaffer et al., 2020) to obtain the functional potential and the genomic taxonomy database (GTDB (Parks et al., 2018, 2020)). For unclassified species, we manually annotated the taxonomy based on the phylogenetic tree constructed using the GTDB marker genes. Species that contain a reference genome of an isolate were counted as cultured, even when they might not be available from official culture collections. Similarly, species named after an isolated strain in GTDB were annotated as cultured.

### CMGM species comprehensively cover the mouse gut metagenome

The CMGM genome collection represents 1’699 species, and we discovered 226 new genera, 7 new families, and 1 new order (Fig. 1A). 78.1% of the CMGM species are uncultured, with only 15% having a mouse-specific cultured strain. 164 species do not have a cultured species even at the order level. The sum of cultured species accounts on average for less than 25% of the mouse metagenome.

**Fig. 1.**
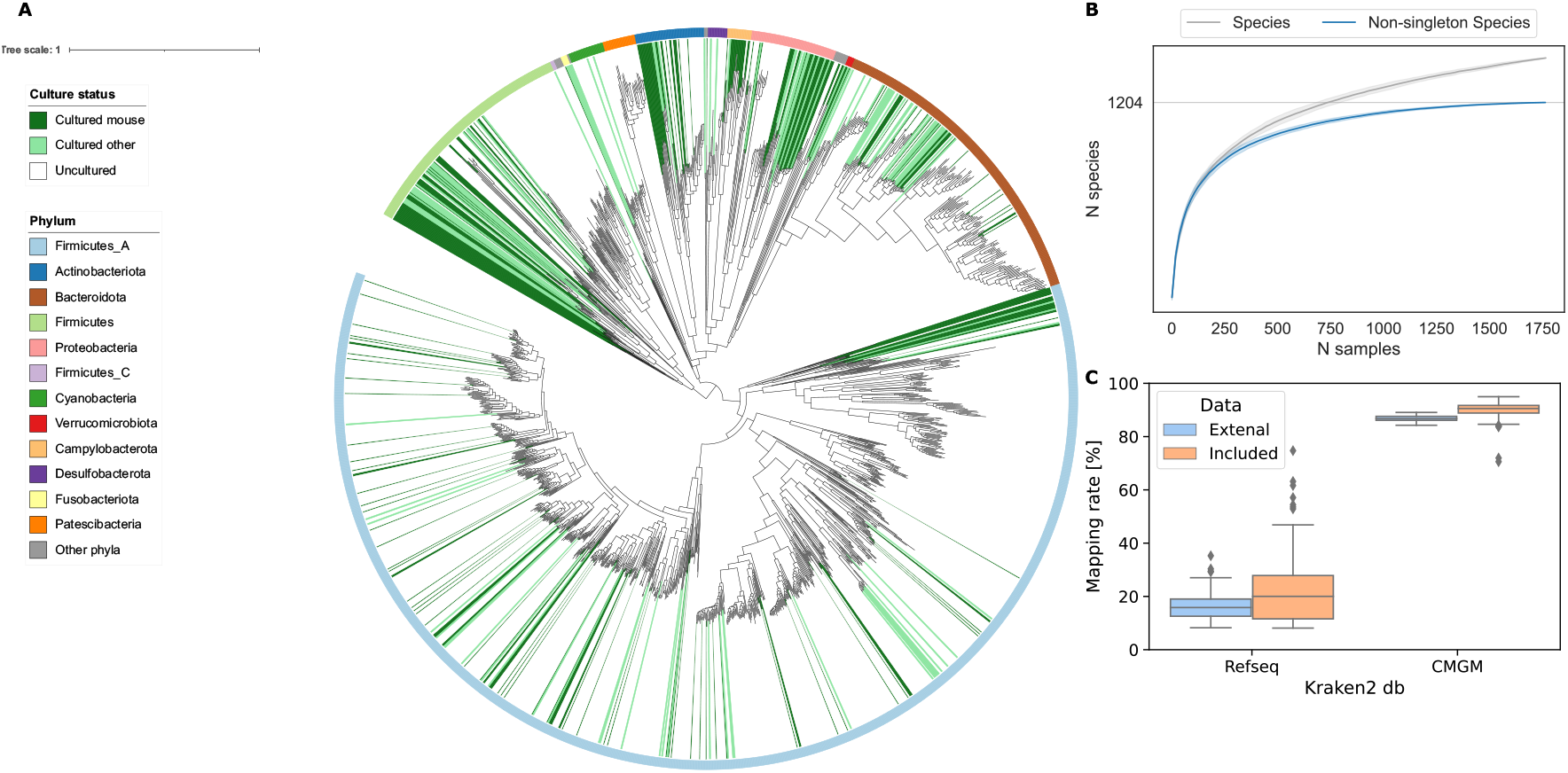
CMGM comprehensively covers the mouse metagenome. **A,** Maximum-likelihood phylogenetic tree of the 1699 bacterial species detected in the mouse gut. Clades are colored by culture status, and the color ring indicates the phylum. **B,** Rarefaction curves of species. **C,** Comparison of mapping rates of the mouse gut metagenome.

**Fig. 2.**
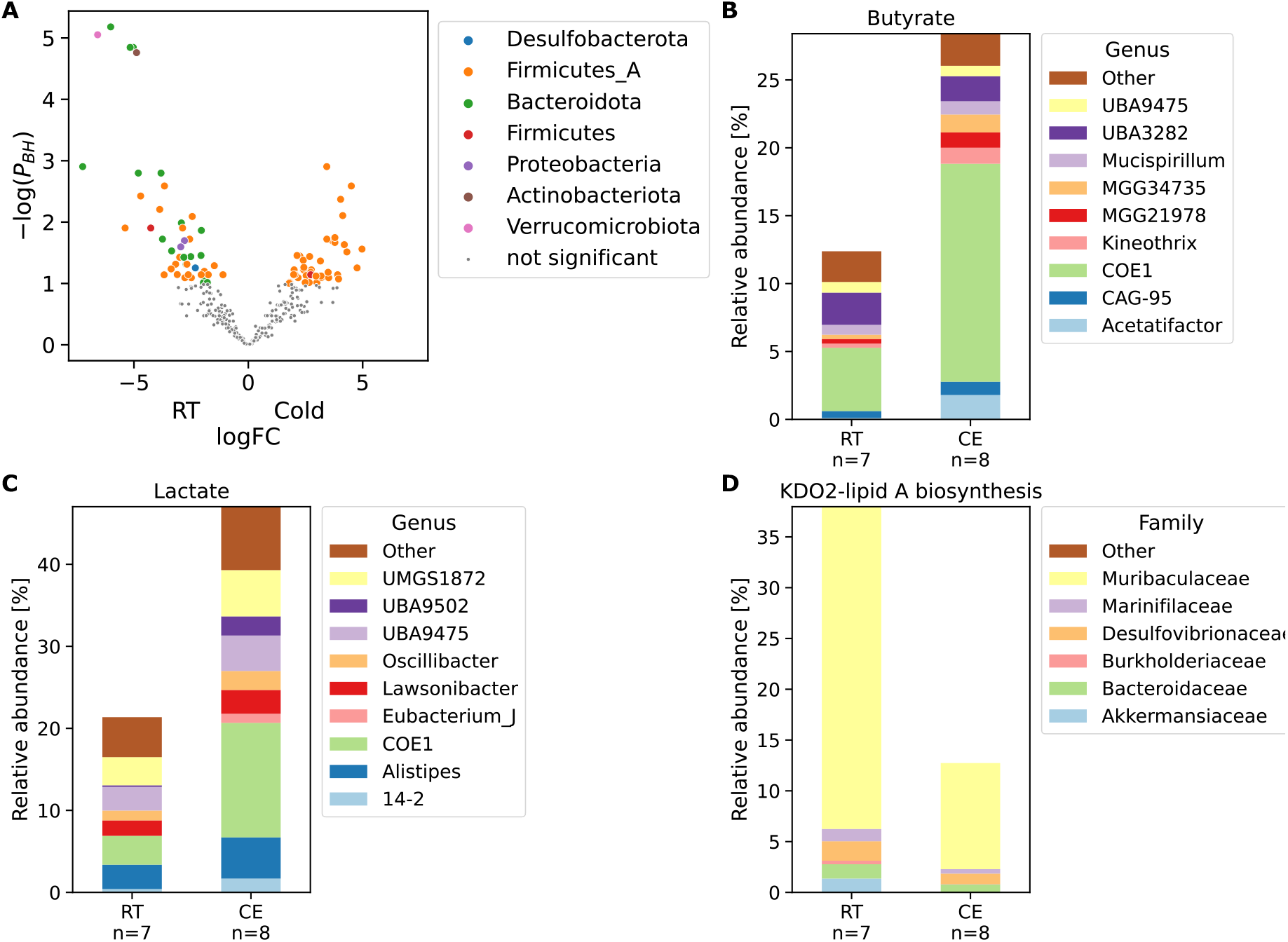
CMGM links functional changes to driver species. **A,** Volcano plot of species changes in mouse cecal microbiota upon cold exposure. Significantly changed species are colored by their phylum. P_BH_: P-value corrected for multiple testing using Benjamini-Hochberg **procedure. B-D,** Bar plots of significantly changed pathways in mouse cecal microbiota upon cold exposure. The contribution to the relative abundance of each module is partitioned by genus B+C and family D. CE: Cold exposure, RT: Room temperature control

Rarefaction analysis shows that the number of species reached a saturation point at 1240 species when considering species with at least two conspecific genomes (Fig. 1B). This indicates that the CMGM catalog contains all species commonly living in the mouse gut. More rare species can still be discovered, as indicated by the non-converging rarefaction curve with singletons (species that were recovered in one only sample). CMGM achieves a mapping rate of the mouse metagenome of 90.5% using Kraken2 (Wood et al., 2019), which is a 4.5-fold increase compared to the standard Kraken database that contains all RefSeq genomes from archaea, bacteria, viruses, and plasmids (Fig. 1C). To independently evaluate the mapping rate of the CMGM catalog, we used an external dataset of cecum samples, which was explicitly left out from this catalog. The CMGM species covered 87.7% of the metagenomic reads, which represents an over 5.5-fold increase to the RefSeq database (Fig. 1C).

### CMGM enables comparative analysis of mouse metagenomes by relating functional changes to driver species

To illustrate how this catalog allows discovering compelling biological insights, we analyzed the metagenome from mice exposed to cold ambient air temperature. Cold exposure is a stimulus that activates the classical brown fat and promotes beige cell development within the subcutaneous white adipose tissue (Cannon and Nedergaard, 2017; Chechi et al., 2013; Stojanović et al., 2018). As such, it is an extensively used intervention for enhancing thermogenic and mitochondrial activity in adipose tissues, leading to decreased adipose tissue amount and improved glycemic status. We (Chevalier et al., 2015), and others (Ziętak et al., 2016) showed that cold exposure leads to a marked shift of the microbiota composition observed by 16S analysis, which is in itself sufficient to improve the insulin sensitivity, induce tolerance to cold, increase the energy expenditure and lower the fat content– an effect in part mediated by activation of the brown fat (Chevalier et al., 2015; Ziętak et al., 2016) and browning of the white fat depots in the cold microbiota-transplanted mice (Chevalier et al., 2015; Cypess et al., 2015; Ghorbani et al., 1997; Guerra et al., 1998; Kopecky et al., 1995). These results indicate an existence of a microbiota-fat signaling axis; however, the signaling cascades mediating this process remain poorly understood.

As noticed previously (Chevalier et al., 2015), here we confirmed that *Akkermansia muciniphila*, the only representative of the phylum *Verrucomicrobiota* was eliminated by cold exposure (Fig. 3A). We found that cold exposure leads to a decrease of the family *Muribaculaceae* and an increase of *Lachnospiraceae* and *Oscillospiraceae.* The species NM07-P-09 sp004793665 (the only species from the phylum *Actinobacteriota*) and three other *Muribaculaceae* species were equally significantly decreased (Fig 3A, P_BH_ < 1e-4). The phylum *Proteobacteriota*, consisting of two species, was significantly decreased. On a functional level, cold exposure led to a doubling of butyrate and lactate production. These changes were mainly due to the increase of the family *Lachnospiraceae,* specifically the increase of the uncultured genus *COE1* (Fig. 3B, C). To address whether these uncovered metagenomic changes are indeed reflected in differences of the actual metabolite levels, we looked at the germ-free mice transplanted with microbiota from the cold-exposed mice or from their RT-kept controls. Transplantation of the cold-adapted microbiota led to an increase in the production of butyrate, lactate, propionate, and succinate in the recipients’ cecum compared to germ-free mice inoculated with microbiota from control mice (Fig. S3). Interestingly, the increased lactate was also measured in the cecum and serum of mice with an intermittent fasting feeding regime (Li et al., 2017), which has been shown to induce browning via the induction of the Vascular endothelial growth factor (Kim et al., 2017). Similarly, succinate is linked to the increase of thermogenesis (Mills et al., 2018). We found a decrease of the prokaryotic succinate dehydrogenase, which metabolizes succinate to fumarate, suggesting a mechanistic link between the cold-induced microbiota changes and the adipose tissue browning.

**Fig. 3.**
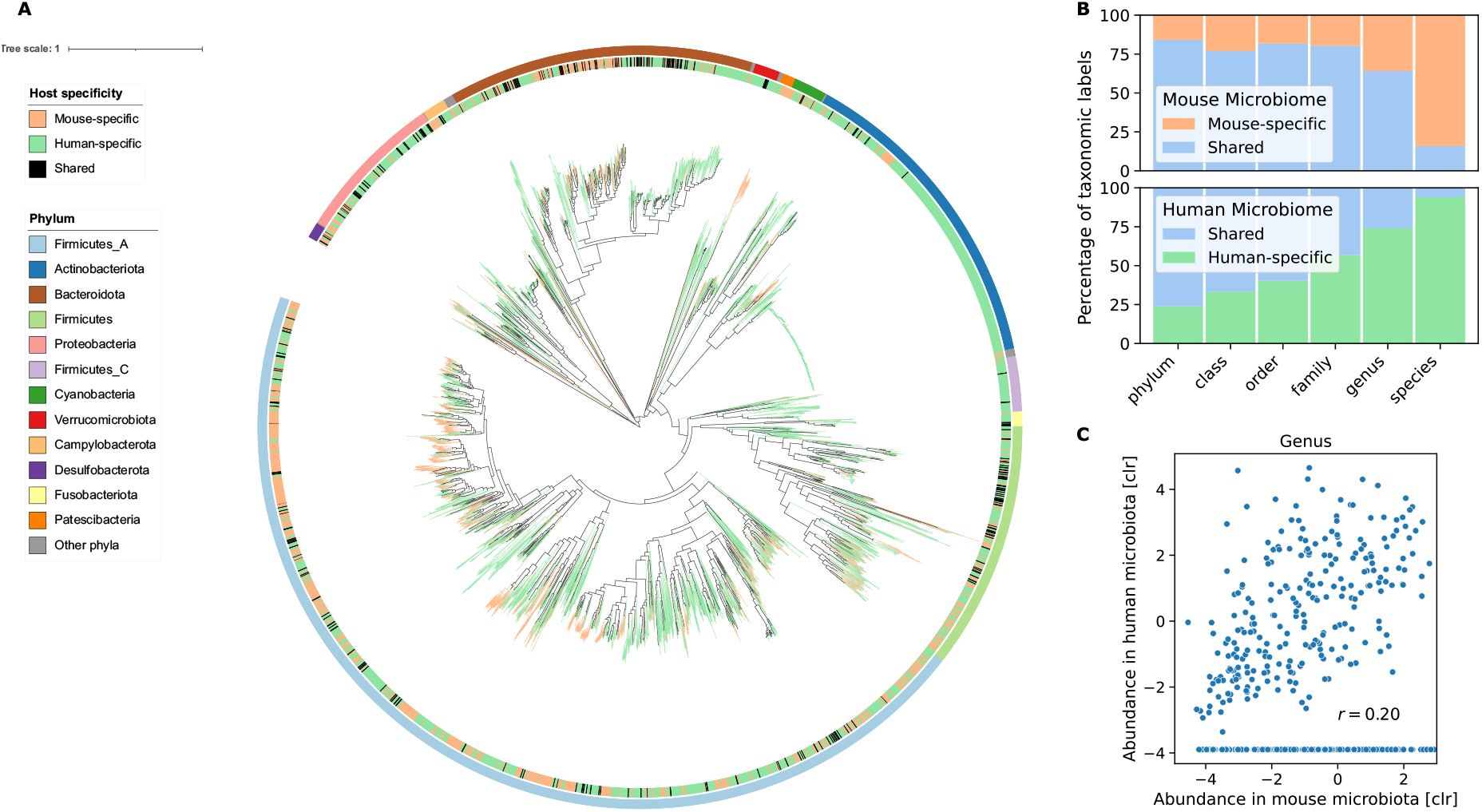
Human and mouse guts harbor distinct bacterial species. **A,** Maximum-likelihood phylogenetic tree of the bacterial species from UHGG and CMGM. The innermost color ring and the tree branches are colored by host specificity. The outermost color ring indicates the phylum attribution. **B,** Percentage of shared and host-specific taxa from CMGM and UHGG at different taxonomic levels. **C,** Correlation of average abundance of genera in human and mice microbiotas. Genera that are not detected in human but present in mouse have an imputed value of −4. CLR = centered log ratio.

We also observed a decrease in Lipopolysaccharide (LPS) synthesis, both in an LpxL-LpxM–dependent and –independent way, primarily attributed to the cold-induced reduction of *Muribaculaceae* (Fig. 3D). LPS administration leads to reduced core body temperature and heat release, correlated with mitochondrial dysfunction (Okla et al., 2015). In contrast, genetic deletion of the LPS receptor, the toll-like receptor 4 (TLR4), leads to resistance against high caloric diet-induced obesity, improved glucose tolerance and insulin sensitivity, and adipose tissue browning (Fabbiano et al., 2018). These findings suggest an additional possible link between the cold-induced microbiota changes and adipose tissues both at mechanistic and bacterial level, contributing to improved insulin sensitivity and browning of the white fat

This example illustrates the CMGM catalog’s usability as a reference for metagenomic studies, enabling discovering precise and comprehensive changes of species and the related function induced by a treatment or a disease. The CMGM sets the ground for reanalysis of the existing datasets for uncovering species and bacterial functions that are involved or altered by the condition of interest.

### Comparison between human and mouse gut microbiomes

Studying mouse microbiota and its impact on the host as a proxy for humans implies their similarities. However, 16S rDNA profiling and gene catalogs do not allow a comprehensive analysis of the analogy between human and mouse microbiota down to species level. Also, much fewer species from the mouse gut are sequenced than from the human gut (Hugenholtz and de Vos, 2018). The CMGM catalog, together with the recent creation of genome collections from the human gut (Almeida et al., 2021), renders this comparison possible. Here, we compared the species from CMGM to the ones from the unified human gastrointestinal genomes (UHGG) (Almeida et al., 2021) and applied the same criteria as for clustering (ANI > 95%). We annotated all species from both hosts with the genome taxonomy database (GTDB, release 5) and curated the unannotated taxonomic levels to allow a consistent taxonomic comparison from domain down to species level.

More than half of the species in both microbiomes belong to the phyla *Firmicutes_A* (Fig. 3A)*. Firmicutes_A* and *Bacteroidota* (*Bacteroidetes*) are the most abundant phyla in both human and mouse microbiomes. Overall, 16 phyla have representatives in both human and mouse microbiome, and 5 are only found in human and not in mice. In contrast, the phyla *Deferribacterota, Thermotogota,* and the two species *Chlamydia muridarum* and *Chlamydophila psittaci,* which represent an own phylum, are specific to mice. No archaea were reconstructed from the mouse gut metagenome, whereas 0.4 % of the genomes in the human gut from the UHGG belong to this domain. At the family level, humans and mice share 98 of the 122 taxa (80% overlap, Fig. 3B), whose average abundance in human and mouse microbiota are correlated (r=0.55, p=1.1e-09). The families *Lachnospiraceae*, *Oscillospiraceae*, and *Butyricicoccaceae*, have high abundance in both human and mice. The family *Muribaculaceae* is 5.6 times more abundant in mice than in humans, whereas *Bacteroidaceae* is 4 times less. While at the genus level, 282 of 440 of taxa are shared (64% overlap, Fig. 3B) in line with results based on 16S rDNA sequencing (Krych et al., 2013, the abundance of the genera showed only a weak correlation (Fig. 3C, r=0.20, p=5e-05). Intriguingly, the genus *Collinsella* (phylum *Actinobacteria*), associated with atherosclerosis and rheumatoid arthritis (Chen et al., 2016; Karlsson et al., 2012), is represented with 579 species in the human but not found in the mouse metagenome.

Strikingly, from the 1699 CMGM species, only 271 (16%) were identified in the human gut microbiota (Fig. 3A, B). The shared species account, on average, for 14.4% of the mouse gut microbiome, and 36 of the 1’699 species in CMGM belong to mouse-specific families. This shows major differences between human and mouse microbiota at species level, demonstrating that mice and human microbiota are largely distinct. These findings effectively challenge our view on the analogy between human and mouse microbiota and may impact the experimental designs, analyses and approaches for studying the human gut microbiome using mouse as a proxy for human.

## Discussion

We generated a comprehensive catalog of the mouse gut metagenome: 50’011 genomes from 1’699 species. This resource enables the mapping of over 90% of the mouse metagenome. Three-quarters of the species are uncultured. Some do not even have a representative at the order level, pointing to the CMGM catalog as a basis for targeted culturing of these missing strains.

Saturation in the rarefaction analysis shows that the CMGM catalog contains all species commonly living in the mouse gut. Nevertheless, we cannot exclude that new samples may add diversity that is not part of the CMGM, for example, species present in single samples or wild mice. However, CMGM is built by assembling all publicly available data from the most used mouse strains, thus comprehensively representing the microbiome of laboratory mice. Comparing the mouse microbiota to the human counterpart reveals overlap and correlation of the average abundance from phylum down to family level. As suggested by amplicon sequencing (Krych et al., 2013), the genera are qualitatively the same but quantitatively different. We observed only a medium correlation between their average abundances in human and mouse microbiota. Whereas a comprehensive and precise comparison at species level between the two microbiomes was not previously feasible (Hugenholtz and de Vos, 2018; Johnson et al., 2019), the comparison of CMGM with the UHGG collection reveals an overlap of only 16% of the species.

While the overlap at the genus and higher taxonomic levels may imply a functional similarity of the human and mouse microbiome, this assumes that functions are conserved within a taxon. While for some functions this is indeed the case, the functional annotation is precisely biased towards conserved functional annotations, which can be transferred from model organisms to less-studied bacterial species. Species from the same genus, even strains from the same species, can have contrasting functions. Strains from the same species can differ in up to 30% in their gene content (Van Rossum et al., 2020) that may help strains from the same species to adapt to different environments. This is especially well studied for the species *Limosilactobacillus reuteri,* which has mouse- and human-adapted strains, however with very different functions (Dheilly et al., 2020; Frese et al., 2011). The different abundance of the mouse and the human microbiome at the genus level, indicted by the weak correlation of average abundance (r = 0.2), compromises even the transferability of the conserved functions at genus level.

Different ways can be envisaged to overcome these challenges. For example, creating ‘humanized’ mouse models by advanced transplantation of human gut microbiota into germ-free mice, or complementing the work by exploring additional animal models (Nguyen et al., 2015). To leverage data produced using conventional mice, it will be important to uncover functional homologs between the species adapted to mouse and human microbiota, e.g., by identifying ‘guilds’ (Root, 1967), groups of species that use the same type of resources in a similar way. The provided consistently functionally annotated species of the human and mouse microbiome lay the basis for such work.

The mouse studies included in the CMGM contain biological insights that were not accessible previously, for example, because they relate to previously unknown species, and incomplete genomes. The knowledge of the genomes and a nearly complete mapping rate is a basis for precise analysis on functional level. This enables uncovering species and bacterial functions that are involved or altered by the condition or treatment of interest. Our resource containing comprehensive collection of the species from the mouse gut and their functional capacity, sets the ground for reanalysis of the existing datasets, and allows analysis of the mouse gut microbiome at an unprecedented depth.

## Author contributions

S.K. wrote the code, analyzed, and interpreted the data, and generated the figures. E.Z. and M.T. guided the project, interpreted the data, and supervised the work. All authors conceptualized the study and wrote the paper.

## Acknowledgments

We are grateful to Christopher Rands for the critical reading of the manuscript and to all members from our labs for discussions. This project has received funding from the European Research Council (ERC) under the European Union’s Horizon 2020 research and innovation programme (ERC Consolidator Grant agreement No. 815962, Healthybiota) to M.T.

## Supplementary Figures

**Fig. S1.**
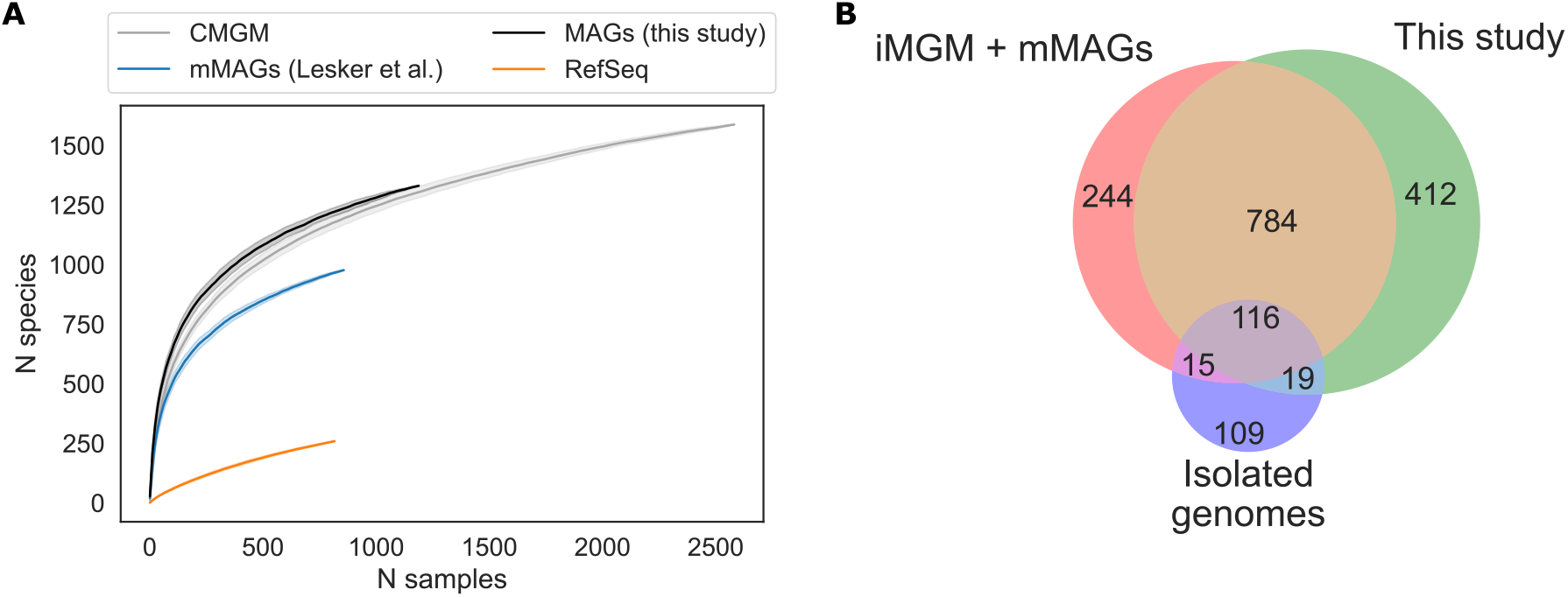
Metagenome assembled genomes radically increase number of species from the mouse gut. **A,** Rarefaction curve of number of species recovered per sample from different sources and CMGM as a whole. **B.** Venn diagram of the species present in the different subsets of CMGM.

**Fig. S2.**
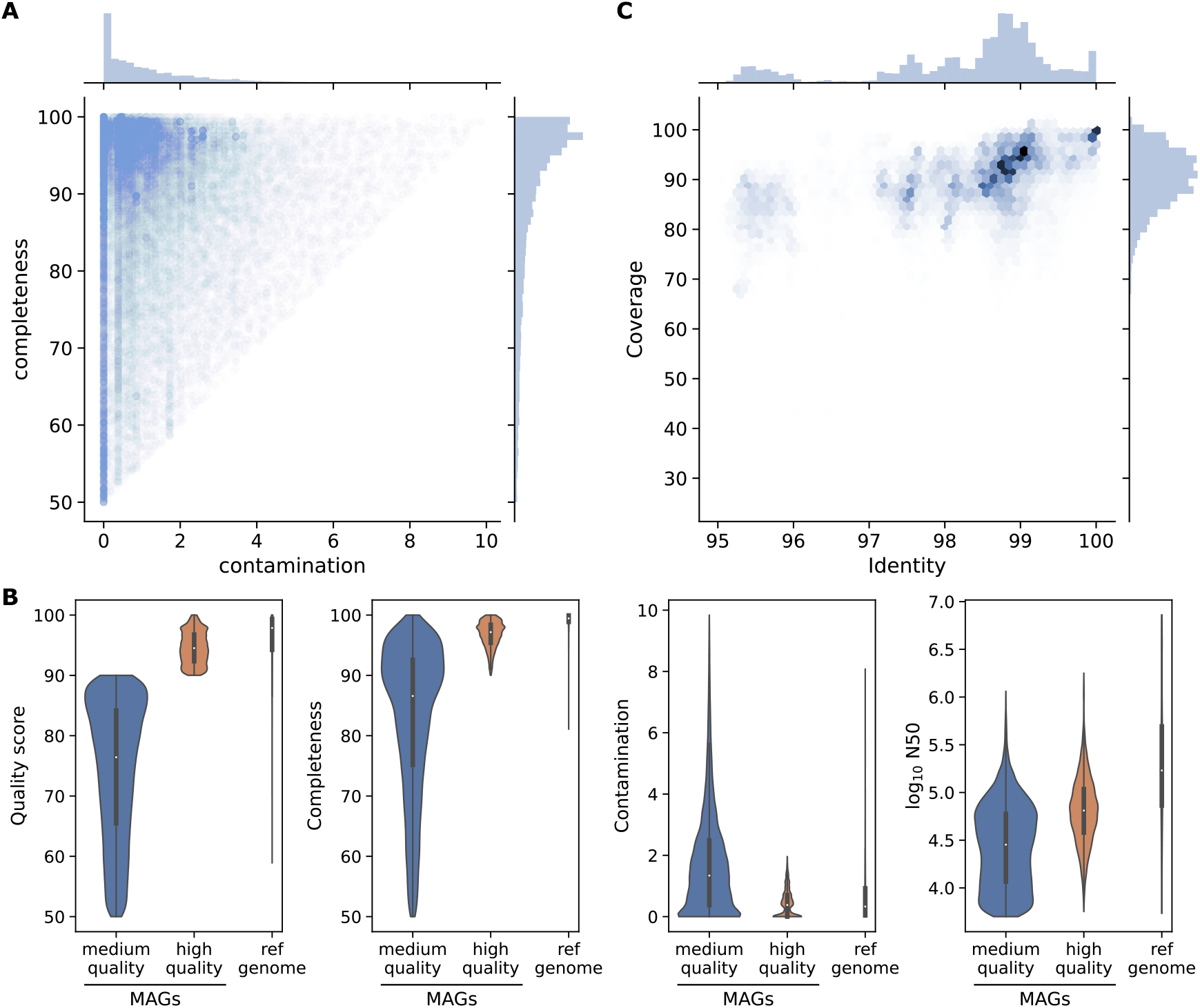
Many metagenome-assembled genomes have comparable quality to reference genomes. **A,** Distribution of the MAGs included in the CMGM collection according to their completeness and contamination estimated with checkM. MAGs with ‘completeness −5×contamination’< 50 were excluded. **B,** Violin plots showing the quality score, completeness, contamination estimated using checkM and the log_10_ N50 from the assembly for the reference genomes and MAGs present in CMGM. **C,** Density plot of the coverage vs. identity of the MAGs alignments to 494 reference genomes

**Fig. S3.**
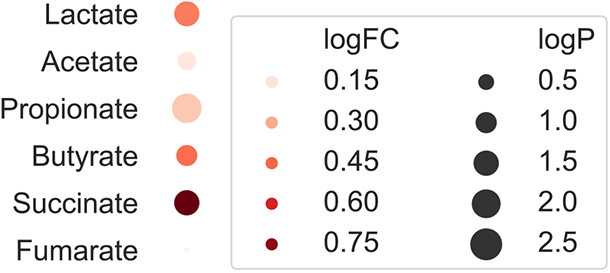
Metabolite changes by cold adapted microbiome. Dot-plot of metabolite changes in ceca of germ-free mice transplanted with cold-adapted microbiota compared to RT-microbiota transplanted controls.

## Supplementary Data

Available from https://ezmeta.unige.ch/CMGM/v1/SuppTables

**Table S1 | Metagenome samples used to construct CMGM.**

The table shows the metagenome samples used for the generation of CMGM. The CMGM_Id corresponds to the SRA read id, except for the samples sequenced by our lab. The table contains information retrieved from NCBI that was available for most of the samples: Name, description, Link to bioproject, collection data, country, and submission center. The column ‘Source’ specifies the organ from which the sample was taken. If the information was available in any of the metadata. Samples of the cold-adapted microbiota under the bioproject accession PRJNA646351 were sequenced for this study.

**Table S2 | Reference genomes associated with the mouse gut.**

The table shows the assembly information of reference genomes associated with the mouse gut. These genomes were filtered for completeness and contamination before integration into CMGM. The columns ‘Isolated’ and ‘Cultured’ label if the genome is Isolated and cultured. The ‘collection’ describes if the genome is part of a mouse-specific culture collection. The genomes of the miBC collection are assembled for this study.

## Methods

### Sequencing of metagenomic data of mice

The sample collection and metagenomic sequencing were approved by the Swiss federal and Geneva cantonal authorities for animal experimentation (Office Vétérinaire Fédéral and Commission Cantonale pour les Expériences sur les animaux de Genève). Animals were on C57Bl/6J background, commercially available through Charles River, France. The mice experiment is detailed in (Chevalier et al., 2015). Paired-end metagenomic libraries were prepared from 100 ng DNA using TruSeq Nano DNA Library Prep Kit (Illumina) and size selected at about 350 bp. The pooled indexed library was sequenced in a HiSeq4000 instrument at the iGE3 facility (University of Geneva).

### Collection of public metagenome and genomic data

We searched the sequence read archive (SRA, accessed December 2019) of the National Center for Biotechnology Information (NCBI) for all publicly available paired-end metagenome runs from the mouse microbiome. We specifically excluded samples from human origin and amplicon sequences and different body parts than the gut. We extracted 1226 metagenome runs belonging to 45 projects. Metadata was retrieved using BioServices (Cokelaer et al., 2013) and curated (Table S1). We retrieved 776 assemblies from RefSeq who were linked to a biosample collected from mouse (Table S2). We excluded reference genomes collected from other body parts than the gut or feces.

### Metagenome assembly and binning

Metagenomics and genomic reads were processed using the metagenome-atlas v2.3 (Kieser et al., 2020) pipeline with the command ‘atlas run genomes’. In short, using tools from the BBmap suite v37.78, reads were quality trimmed, and contaminations from the mouse genome were filtered out. Reads were error corrected and merged before they were assembled with metaSpades v3.13 (Nurk et al., 2017). Contigs were binned using metabat2 v 2.14 (Kang et al., 2019) and maxbin2 v2.2 (Wu et al., 2016), and their predictions were combined using DAS Tool v 1.1.1 (Sieber et al., 2018). For the assembly of the 53 genomes of the mouse intestinal bacterial collection, we used the assembly workflow of metagenome-atlas and set ‘spades_preset: normal’ which uses the basic spades as assembler. The quality of the genomes was estimated using checkM v1.1 (Parks et al., 2015).

### Genome filtering and species clustering

Genomes with good enough quality were grouped into species with average nucleotide identity (ANI) > 95%. First, genomes with an estimated quality of ‘completeness-5*contamination’ < 50 or N50 <= 5000 were excluded. Then, all pair-wise average nucleotide identities (ANI) above 0.8 were calculated using bindash (Zhao, 2018). The genomes were pre-clustered into clusters that contain at least one pair of genomes above the threshold. Then each cluster was grouped into species by hierarchical clustering with average linkage using scipy (Virtanen et al., 2020). For each species cluster, the genome with the highest score based on the following formula was selected as the representative.

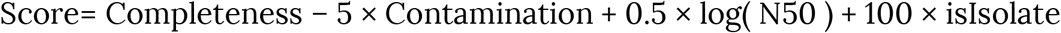

Where Completeness and Contamination are estimated using checkM v1.1 (Parks et al., 2015), N50 is the N50 score of the assembly contiguity, and ‘isIsolate’ is 1 for isolates and 0 for MAGs, to ensure that isolated genomes are preferred over MAGs even if they have lower quality.

### Phylogenetic and taxonomic analysis

The species representatives of both the CMGM and the unified human gastrointestinal genomes (UHGG) (Almeida et al., 2021) were annotated using the genomic taxonomy database toolkit (GTDB-tk v1.2 (Parks et al., 2018)) and the GTDB release 5. A maximum-likelihood tree for the CMGM alone and the CMGM combined with the UHGG based on the 120 bacterial marker genes from the GTDB was built using fasttree v2.1 (Price et al., 2010) and rooted at the midpoint. The phylogenetic trees are visualized with iTOL v5 (Letunic and Bork, 2019).

Unannotated, taxa from both human and mouse were manually annotated. New orders, families and genera were defined at the relative evolutionary divergence of close related annotated taxa.

### Inferring cultured status

Species that contain a reference genome included in the CMGM catalog are counted as cultured from a mouse origin. If GTDB-tk (Parks et al., 2018) was able to annotate the species to a reference with ANI >95%, we counted the species as cultured from a non-murine source. In both cases, if the reference genome was excluded from RefSeq (i.e., metagenome-assembled genomes) or labeled as uncultured we counted the species as isolated but not cultured.

### Quantification

We build Kraken 2 and bracken (Wood et al., 2019) databases for the CMGM and the UHGG based on our curated taxonomy. The mapping rates were calculated as a fraction of the reads attributed with bracken at the species level to the total reads. For comparison, we quantified reads using the standard Kraken2 database accessible from https://benlangmead.github.io/aws-indexes/k2 (stand December 2020). For most quantification, the mapped reads per genome were summed, and the centered log-ratio (CLR) was calculated using the sci-kit bio package (http://scikit-bio.org/) after imputing zeros using a multiplicative replacement approach. To calculate the average species abundance in the mouse and human metagenome, we used 184 fecal samples from the MGC v1 (Xiao et al., 2015) and a random subset of 1000 samples of the human metagenome that is commonly used for benchmarking (Almeida et al., 2019). The Pearson correlation between the abundance of taxonomic groups in the human and mouse microbiota was performed with scipy v1.4.1 (Virtanen et al., 2020). When relative abundance was used as a measure, we used BBsplit (https://jgi.doe.gov/data-and-tools/bbtools/bb-tools-user-guide/) with the parameters’ ambiguous2=best minid=0.9’ to map metagenomic reads to the references with 90 identity. We estimated the genome coverage as the median of coverage over 1000bp blocks.

### Functional annotation

The species representatives of both the CMGM and the UHGG were annotated using DRAM (Shaffer et al., 2020). A Kegg-module is inferred to be present if ¾ of all the steps were present in a genome. As there are no modules for short chain fatty acids in Kegg we created custom modules (see the ‘Code’ section). The step-coverage was calculated with DRAM for all Kegg-modules. The metagenome-side abundance of functional modules was calculated as the sum of the relative abundances of all genomes containing a module. We used the Welch test and Benjamini-Hochberg correction to estimate the significance of changes in module abundance between experimental groups.

## Data Availability

The metagenomic samples sequenced for this study are available from the NCBI sequence read archive under the project id PRJNA646351. The genomes assembled in this study will be deposited under study accession PRJNA646353 and are accessible now from https://ezmeta.unige.ch/CMGM/v1. The taxonomy, functional annotation, and Kraken databases for both human and mouse are available from the same link.

## Code Availability

The code for the analysis of the cold-exposed microbiota is available from https://github.com/SilasK/CMGM

